# *Methanobacterium nebraskense* sp. nov., a hydrogenotrophic methanogen isolated from saline wetland soil

**DOI:** 10.1101/2025.11.18.688885

**Authors:** Nicole A. Fiore, You Zhou, Karrie A. Weber

## Abstract

A hydrogenotrophic methanogen, designated strain ACI-7^T^, was isolated from an alkaline, saline wetland located in eastern Nebraska, USA. Cells were identified as non-motile rods (0.9–4.2 μm in length and 0.2–0.4 µm in diameter) that occurred singly, in chains, and as long filaments and twisted aggregates. Strain ACI-7^T^ utilized H_2_ + CO_2_ or formate as methanogenic substrates and required growth factors present in yeast extract for continuous cultivation. The strain grew at 20–45°C (optimum, 40°C), at pH 6.5–8.5 (optimum, pH 7.3) and with 0–2.5% NaCl (optimum, 0–1%). The genomic G+C content of ACI-7^T^ was 31.79 mol%. Phylogenetic analysis of the 16S rRNA gene sequence indicated strain ACI-7^T^ was affiliated with the genus *Methanobacterium,* most closely related to *Methanobacterium oryzae* FPi^T^ (97.1% sequence similarity) and *Methanobacterium veterum* MK4^T^ (96.9% sequence similarity). Overall genome relatedness indices for strain ACI-7^T^ compared to other *Methanobacterium* species ranged from 68.45–78.17% for average nucleotide identity and 18.9–25.7% for digital DNA–DNA hybridization. Morphological, physiological, and genomic characteristics indicate that strain ACI-7^T^ represents a novel species, for which the name *Methanobacterium nebraskense* sp. nov. is proposed. The type strain is ACI-7^T^ (=DSM 118696^T^=ATCC TSD-487^T^).

## Introduction

Hydrogenotrophic methanogenesis is an evolutionarily ancient (Berghuis et al., 2019; Ueno et al., 2006) microbial metabolism by which carbon dioxide is reduced to methane using hydrogen as an electron donor. Of the known methanogenic pathways, hydrogenotrophic methanogenesis is also the most widespread, distributed across several taxonomic phyla within the domain Archaea (Berghuis et al., 2019; Buan, 2018). Some of the earliest hydrogenotrophic methanogens studied were associated with the genus *Methanobacterium* (Barker, 1936; Bryant et al., 1967; Paynter & Hungate, 1968; Schnellen, 1947; Smith & Hungate, 1958), which currently includes 27 validly described species (Parte et al., 2020). *Methanobacterium* species have been isolated from several sites, including terrestrial (Krivushin et al., 2010; Shcherbakova et al., 2011), marine (Shlimon et al., 2004), and subsurface (Jain et al., 1987; Kotelnikova et al., 1998; Schirmack et al., 2014) environments. Five of the validly named *Methanobacterium* species have been isolated from wetland ecosystems: *Methanobacterium paludis* (Cadillo-Quiroz et al., 2014) and *Methanobacterium palustre* (Zellner et al., 1988) from peatlands, *Methanobacterium uliginosium* (König, 1984) from a marsh, and *Methanobacterium kanagiense* (Kitamura et al., 2011) and *Methanobacterium oryzae* (Joulian et al., 2000) from rice fields. In this study, we describe a novel *Methanobacterium* species, strain ACI-7^T^, isolated from an alkaline, saline wetland soil located in eastern Nebraska.

### Enrichment and isolation

Wetland soil samples were collected in June 2013 from an alkaline, saline wetland in Lincoln, NE (40.8786, −96.6766) as described previously (Fiore et al., 2025). Initial enrichments were performed with 50% w/v wetland soil in NSW culture medium amended with calcium carbonate as a sole source of inorganic carbon and hydrogen as an electron donor (Fiore et al., 2025). Genome-resolved metagenomic analysis of the initial enrichment community identified five bacterial taxa and one methanogen (Fiore et al., 2025). Metabolic pathways and antibiotic resistance genes identified in the metagenomic dataset were used to develop culture conditions that would select for the methanogen while suppressing growth of the bacterial community members. The methanogen was subsequently enriched from the initial community using MS Enrichment Medium (Bonin & Boone, 2006), prepared and dispensed under Ar/CO_2_ (80:20 v/v) using Hungate technique (Balch & Wolfe, 1976; Hungate & Macy, 1973; Miller & Wolin, 1974) and autoclaved at 121°C for 20 minutes. After autoclaving, sodium sulfide (1 mM), sodium mercaptoethanesulfonate (coenzyme M; 3 mM), and antibiotics (100 µg/mL kanamycin, 100 µg/mL clindamycin, 100 µg/mL tetracycline, and 250 µg/mL vancomycin) were added to the medium from sterile anoxic stock solutions, with a final medium pH of 7.3. After inoculation (10% v/v), hydrogen gas was added via syringe (20 mL H_2_ overpressure for 10 mL culture in 26 mL Balch tubes). Cultures were incubated statically at 37°C in the dark, positioned horizontally to maximize surface area for gas exchange.

Pure cultures were obtained by isolating single colonies from agar shake tubes (Macy et al., 1972) in an anaerobic chamber. One of these cultures was given the strain designation ACI-7 and characterized further. Antibiotics were omitted for subsequent transfers in liquid medium (5% v/v), including those used to evaluate culture purity. Culture purity was confirmed based on microscopic observation, absence of growth with methanogenic substrates omitted, and failed PCR amplification using high coverage, bacteria-specific 16S rRNA primer pairs S-D-Bact-0008-a-S-16/S-D-Bact-1492-a-A-16 and S-D-Bact-0341-b-S-17/S-D-Bact-1061-a-A-17 (Klindworth et al., 2013) relative to positive controls. Pure cultures were stored in at −80°C in anoxic 10% (v/v) glycerol for long-term preservation.

### Morphology and cellular observations

Cells for microscopy were grown to mid-log phase on MS Enrichment Medium with H_2_ and CO_2_ as substrates. Cell morphology, size, motility, and arrangement were observed using a combination of phase contrast microscopy, differential interference contrast microscopy, fluorescence microscopy, and scanning electron microscopy. Phase-contrast microscopy was performed on unfixed samples using an Axioskop 40 FL microscope (Zeiss) equipped with an AxioCam ERc 5s (Zeiss). Differential interference contrast (DIC) microscopy and fluorescence microscopy were conducted using a Nikon A1R-Ti2 confocal laser scanning microscope on cells fixed with 2.5% glutaraldehyde in 0.1 M cacodylate buffer, rinsed twice with the same buffer and once with 10% ethanol, then stained with 10 µM SYTOX Green. Cell length and diameter were estimated from phase contrast and fluorescence micrographs using ImageJ2. F_420_ autofluorescence (Doddema & Vogels, 1978) was examined with an Axioskop 40 FL microscope equipped with an HBO 50 fluorescence illuminator (Zeiss) and an F420 filter set (417/60 nm excitation, 482/35 nm emission, 458 nm beam splitter; AVR Optics). For scanning electron microscopy, cells were fixed with 2.5% glutaraldehyde in 0.1 M cacodylate buffer, ethanol dehydrated, air dried overnight, then sputter coated with a thin layer (∼2 nm) of chromium. The coated samples were examined under a field-emission scanning electron microscope (S-4700, Hitachi). Susceptibility to lysis by sodium dodecyl sulfate (SDS) and distilled water were determined as described by Boone & Whitman (Boone & Whitman, 1988), with cell lysis assessed via optical density and phase-contrast microscopy.

Cells of strain ACI-7^T^ are short rods, 0.9–4.2 µm in length and 0.2–0.4 µm in diameter, occurring as single cells, chains, long filaments, and aggregates (Fig. 1a, Fig. S1). Cells in chains vary from straight to curved, with curved cells often forming helical structures. Some chains form twisted aggregates, either as single chains that twist upon themselves (forming a loop at one end) or multiple chains twisted together (Fig. 1b). Similar morphological observations have been described for other taxa within Methanobacteriaceae, including *Methanobacterium aggregans* (Kern et al., 2015) and *Methanothermobacter thermautotrophicus* (formerly *Methanobacterium thermoautotrophicum*) (Zeikus & Wolfe, 1973). Cells are non-motile and stain Gram-negative. Strong autofluorescence is observed at 420 nm (Fig. S2) but rapidly fades. Cells resist lysis by both 0.1% and 1% SDS and by hypotonic solution (distilled water).

**Fig. 1.**
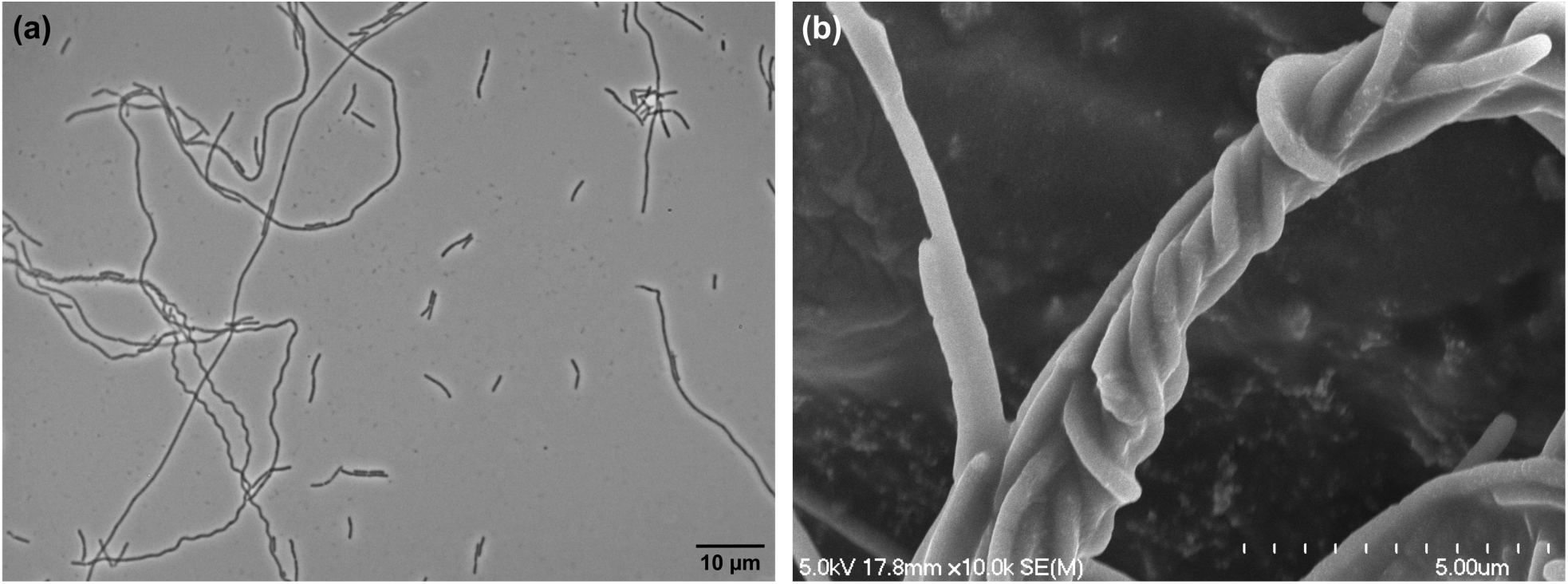
Cell morphology of *Methanobacterium nebraskense* strain ACI-7^T^ as observed with phase contrast microscopy (a) and scanning electron microscopy (b). Cells were grown on H_2_ + CO_2_ under optimal conditions. Scale bars represent 10 µm (a) and 5 µm (b).

### Substrate utilization and growth conditions

Growth and substrate utilization were assessed by culturing strain ACI-7^T^ in MS Enrichment Medium on H_2_ + CO_2_ at 37°C as described above, unless otherwise noted. All tests were performed in triplicate, and growth was monitored by measuring the concentration of methane in the headspace (Powell, 1983). Methane was measured by a gas chromatography with a Varian 430-GC and thermal conductivity detector using ultra high-purity argon as carrier gas, with oven and detector temperatures of 65°C and 250°C, respectively.

The effect of temperature on growth was tested at 4, 20, 30, 37, 40, 45, and 50°C. Incubations from 30–50°C were accomplished with separate incubators at their respective temperatures, 20°C on the bench at room temperature, and 4°C in a laboratory refrigerator. Growth of strain ACI-7^T^ was observed from 20–45°C (optimum 40°C), with no growth at 4°C or 50°C (Fig. S3a). The effect of NaCl on growth was tested with 0%, 1%, 2.5%, 3%, 3.5%, 4.5%, 5%, and 10% NaCl (w/v). Medium was prepared with 0% and 10% NaCl and mixed prior to dispensing to achieve intermediate concentrations. Growth was observed with NaCl concentrations from 0–2.5%, with optimum growth from 0–1% and no growth observed at 3% NaCl (Fig. S3b). The effect of pH on growth was measured at pH values of 6.0, 6.5, 6.9, 7.3, 7.5, 8.0, 8.3, 8.5, and 9.0. The pH of MS Enrichment Medium as described above is 7.3; to achieve other values, the medium was supplemented with organic buffers, as recommended for bicarbonate-buffered media (Prakash et al., 2023). The organic buffers used were as follows, all at 50 mM concentrations: pH 6.0 (MES), pH 6.5–6.9 (PIPES), pH 7.5–8.5 (HEPES), and pH 9.0 (CHES). Adjustments to pH were made with sterile, anoxic stock solutions of 1 M HCl or 1 M Na_2_CO_3_ prior to inoculation. pH was measured immediately after inoculation and increased by no more than 0.2 units after 5 days of incubation. Growth of strain ACI-7^T^ was observed over pH 6.5–8.5 (optimum 7.3), with no growth at pH 6.0 or 9.0 (Fig. S3c).

To determine nutrient requirements, all sources of organic carbon (yeast, tryptone, and coenzyme M) were omitted from the medium. Growth of strain ACI-7^T^ was tested in minimal medium alone and with the following additions: yeast extract (0.5 g L^-1^), tryptone (0.5 g L^-1^), casamino acids (0.5 g L^-1^), acetate (10 mM), coenzyme M (3 mM), and Wolfe’s vitamin solution (Wolin et al., 1963). Growth was monitored for six successive transfers at 5% (v/v) inoculum. An unidentified component of yeast extract was required for stable cultivation of strain ACI-7^T^; casamino acids, acetate, and vitamins were not adequate substitutes. Coenzyme M alone was sufficient for four successive transfers, but methane production declined with the fifth and sixth transfer.

The following compounds were tested as potential substrates for methanogenesis: H_2_ + CO_2_, formate, acetate, dimethylsulfide, methanol, 2-propanol + CO_2_, H_2_ + methanol, H_2_ + methylamine, methylamine, dimethylamine, and trimethylamine. All aqueous substrates were tested at concentrations of 10 mM, with positive results confirmed by three successive transfers and compared to no-substrate controls. When testing growth on substrates other than carbon dioxide in combination with hydrogen (e.g. H_2_ + methanol, H_2_ + methylamine), the bicarbonate buffer was replaced with 50 mM HEPES and medium was prepared and dispensed under 100% Ar headspace. H_2_ + CO_2_ and formate were the only substrates used for growth and methanogenesis, with specific growth rates of 0.033 hr^-1^ and 0.016 hr^-1^ under optimal conditions, respectively. Acetate, dimethylsulfide, methanol, 2-propanol + CO_2_, H_2_ + methanol, H_2_ + methylamine, methylamine, dimethylamine, or trimethylamine were not used as substrates for methane production.

### Genome and phylogenetic classification

The genome sequence for strain ACI-7^T^ was obtained as previously described (Fiore & Weber, 2025). Briefly, DNA was extracted using a method modified from Zhou *et al*. (Zhou et al., 1996), and library preparation and sequencing (PacBio Sequel IIe) were completed by the University of Delaware DNA Sequencing and Genotyping Center. Reads were assembled with Flye 2.9.1 (Kolmogorov et al., 2019), resulting in a single, circular contig 2,239,949 bp in length at an estimated 5253 × coverage with a G+C content of 31.79 mol% (Fiore & Weber, 2025).The genome was 99.2% complete with no contamination, based on conserved single-copy genes for Euryarchaeota using CheckM v1.1.2 (Parks et al., 2015). Genome annotation with the NCBI Prokaryotic Genome Annotation Pipeline (PGAP) 6.8 (Tatusova et al., 2016) predicted 2,411 genes representing 2,360 protein coding sequences, 2 complete sets of rRNAs (5S, 16S, and 23S rRNAs), 37 tRNAs, and 2 noncoding RNAs (ncRNAs). The two 16S rRNA gene sequences were extracted from the genome assembly using Barrnap 0.9 (https://github.com/tseemann/barrnap), both of which were 1478 bp in length with 100% identity. Additional annotation was performed with BlastKOALA against the KEGG GENES database (Kanehisa et al., 2016). Genomic evidence for hydrogenotrophic methanogenesis in ACI-7^T^ includes all necessary genes for the Wolfe cycle of carbon dioxide reduction to methane (Thauer, 2012) and the archaeal type Wood-Ljungdahl pathway of carbon fixation (Borrel et al., 2016) (Table S2).

To confirm the authenticity of the genome assembly, a partial 16S rRNA gene sequence was obtained via Sanger sequencing for comparison to the genome-derived sequences (Chun et al., 2018). The 16S rRNA gene was PCR amplified from strain ACI-7^T^ using the primers A109F (Whitehead & Cotta, 1999) and Met1340R (Wright & Pimm, 2003) and sequenced by Eurofins Genomics (Louisville, KY), resulting in a partial 16S rRNA gene sequence 1045 bp in length. The 1045 bp 16S rRNA gene sequence obtained via Sanger sequencing was identical to the full-length gene sequence obtained from the genome, with no gaps or mismatches. The full length 1478 bp 16S rRNA gene sequence extracted from the genome assembly was used for phylogenetic analysis. Both phylogenetic (16S rRNA) and phylogenomic analyses were used to infer the taxonomy and phylogeny of strain ACI-7^T^. Phylogenetic analysis based on the 16S rRNA gene was performed for strain ACI-7^T^ with the type strain of all validly published species in the genus *Methanobacterium* (Table S1). The 16S rRNA sequences were obtained from GenBank and aligned with MUSCLE 5.1 (Edgar, 2022). The alignment was trimmed with BMGE (Criscuolo & Gribaldo, 2010) and maximum-likelihood phylogenetic trees were inferred using RAxML v8.2.12 (Stamatakis, 2014) using 1000 bootstrap replicates. Pairwise sequence similarities were estimated using the EzBioCloud server (Chalita et al., 2024). At the genomic level, overall genome relatedness indices (OGRIs) and phylogenomic trees were determined for strain ACI-7^T^ against all valid *Methanobacterium* spp. with available reference genomes (Table S1). Genomes were obtained from NCBI RefSeq or JGI IMG, with preference for type strains when available (Table S1). For overall genome relatedness, average nucleotide identity (ANI) and digital DNA–DNA hybridization (dDDH) were calculated using OrthoANIu (Yoon et al., 2017) and the Genome-to-Genome Distance Calculator (GGDC) 3.0 with Formula 2 (Meier-Kolthoff et al., 2021), respectively. Maximum-likelihood phylogenomic trees were reconstructed based on 53 conserved archaeal marker genes obtained from the Genome Taxonomy Database (GTDB) R226 (Parks et al., 2020) using PhyloPhlAn 3.1 (Asnicar et al., 2020) and RAxML-NG v1.2.0 (Kozlov et al., 2019) with 100 bootstrap replicates.

Phylogenetic trees reconstructed from 16S rRNA gene sequences indicated strain ACI-7^T^ falls within the genus *Methanobacterium*, forming a cluster including *Methanobacterium oryzae* FPi^T^ and a clade containing *Methanobacterium uliginosum* P2S^T^, *Methanobacterium bryantii* M.o.H.^T^, *Methanobacterium arcticum* M2^T^, *Methanobacterium espanolae* GP9^T^, *Methanobacterium ivanovii* OCM 140^T^, and *Methanobacterium veterum* MK4^T^ (Fig. 2). Based on 16S rRNA gene similarity, strain ACI-7^T^ was most closely related to *Methanobacterium oryzae* FPi^T^ (97.09% sequence similarity), followed by *Methanobacterium veterum* MK4^T^ (96.95% sequence similarity), *Methanobacterium ivanovii* OCM 140^T^ (96.86% sequence similarity), and *Methanobacterium movilense* MC-20^T^ (96.73% sequence similarity), all of which fall below the recommended species-level cutoff of 98.65% (Chun et al., 2018; Kim et al., 2014). At the phylogenomic level, strain ACI-7^T^ formed a highly supported cluster with the clade containing *Methanobacterium bryantii* M.o.H.^T^, *Methanobacterium arcticum* M2^T^, and *Methanobacterium veterum* MK4^T^ (Fig. 3), but not with *Methanobacterium oryzae* FPi^T^ as shown in the 16S rRNA gene tree. *Methanobacterium oryzae* FPi^T^, the closest relative to strain ACI-7^T^ based on 16S rRNA sequence similarity, also had the highest ANI. Compared with strain ACI-7^T^, *Methanobacterium oryzae* FPi^T^ had an ANI value of 78.17%, followed by *Methanobacterium arcticum* M2^T^ and *Methanobacterium bryantii* M.o.H.^T^ with values of 75.56% and 75.46%, respectively (Table S1). Values for dDDH were < 30% for all comparisons against strain ACI-7^T^ (Table S1). Both measures of OGRI fall well below the proposed species boundaries of 95–96% for ANI and 70% for dDDH (Chun et al., 2018; Meier-Kolthoff et al., 2013; Richter & Rosselló-Móra, 2009).

**Fig. 2.**
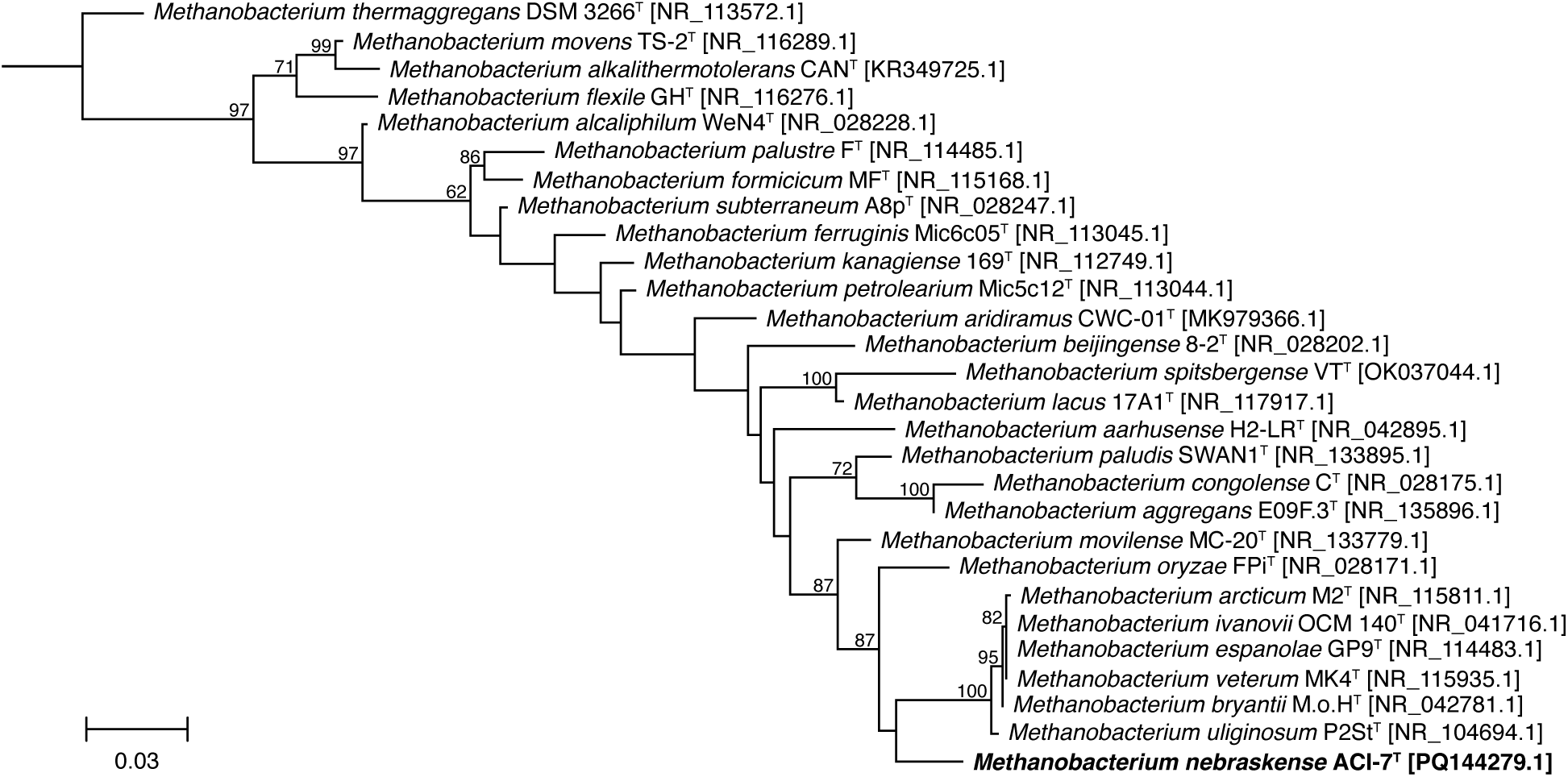
Maximum likelihood phylogenetic tree based on 16S rRNA gene sequences (1326 bp) showing the position of strain ACI-7^T^ and type strains of the genus *Methanobacterium*. Numbers at nodes indicate bootstrap percentages based on 1000 replicates; only values ≥ 50% are shown. Accession numbers are indicated in brackets. *Methanothermus fervidus* V24S^T^ [NR102926.1] was used as an outgroup (not shown). Bar, 0.03 substitutions per nucleotide position.

**Fig. 3.**
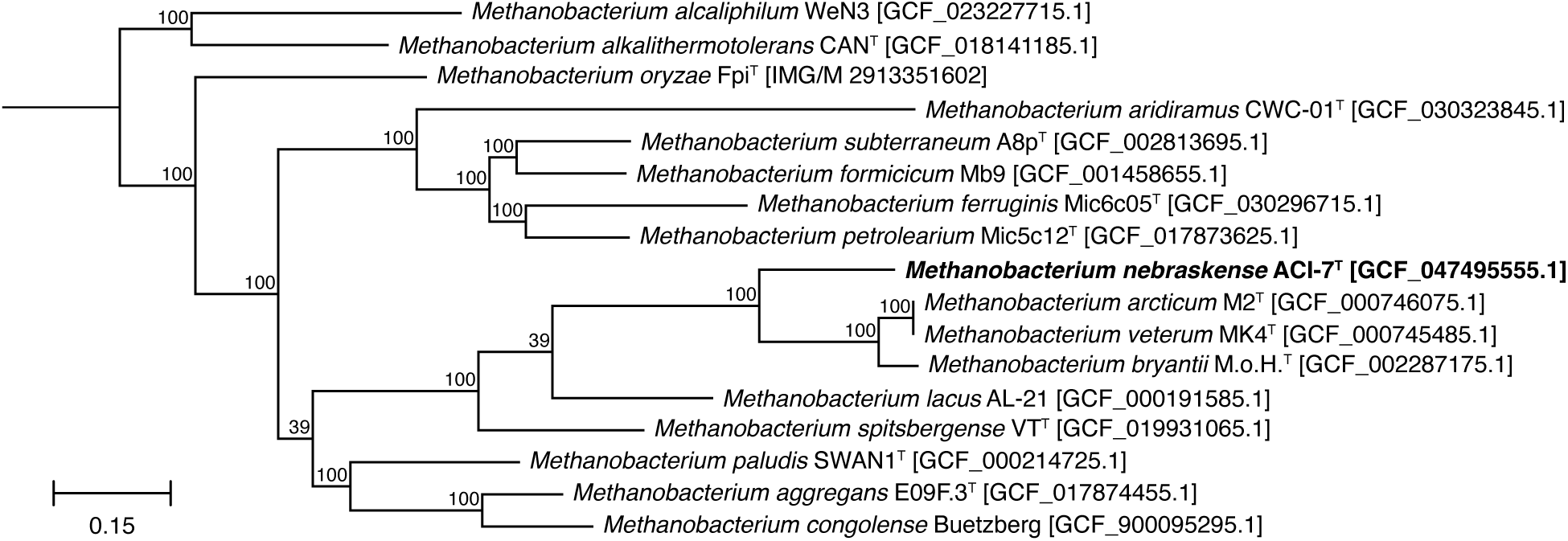
Maximum likelihood phylogenomic tree of strain ACI-7^T^ in relation to other validly published *Methanobacterium* species with available reference genomes. Trees were inferred based on the 53 conserved archaeal marker genes in the Genome Taxonomy Database (GTDB). Values at nodes indicate bootstrap percentages based on 100 replicates. Accession numbers are indicated in brackets. *Methanothermus fervidus* V24S^T^ [GCF_000166095.1] was used as an outgroup (not shown). Bar, 0.15 substitutions per nucleotide position.

The physiological characteristics differentiating strain ACI-7^T^ from the most phylogenetically related type species of *Methanobacterium* are summarized in Table 1. The isolate shared a similar temperature range and optimum with *M. oryzae*, *M. uliginosum*, and *M. arcticum*, but differed considerably from *M. veterum* and *M. movilense*, which grow optimally at lower temperatures. All species including ACI-7^T^ were able to use H_2_ + CO_2_ as substrates, as is characteristic of the genus, but only ACI-7^T^, *M. oryzae*, *M. movilense*, and *M. arcticum* are able to use formate. All species were neutrophilic, but strain ACI-7^T^ and *M. ivanovii* are the most restricted in pH range for growth. Strain ACI-7^T^ tolerated similar NaCl concentrations as *M. oryzae*, both of which were more halotolerant than the other related species. Morphologically, cells of strain ACI-7^T^ were most similar to *M. uliginosum* in size, but formed filaments and aggregates as described for *M. oryzae* and *M. bryantii*. The formation of twisted chains by strain ACI-7^T^ (Fig. 1b) is a distinct morphological trait not documented among its closest relatives. Despite overall physiological similarity between strain ACI-7^T^ and its closest relative, *M. oryzae*, the two species are clearly differentiated by 16S rRNA gene similarity (97.09%), ANI (78.17%), and dDDH (22.0%). The most notable phenotypic differences between ACI-7^T^ and its other most highly related taxonomic relatives are its pH and temperature range as compared to *Methanobacterium veterum* MK4^T^ (96.95% 16S rRNA, 75.33% ANI, 21.1% dDDH) and *Methanobacterium arcticum* M2^T^ (96.47% 16S rRNA, 75.56% ANI, 21.1% dDDH) and ability to use formate as a substrate compared to *Methanobacterium bryantii* M.o.H.^T^ (96.68% 16S rRNA, 75.46% ANI, 21.1% dDDH) and *Methanobacterium veterum* MK4^T^. Based on phenotypic differences, together with genomic and phylogenetic characteristics, strain ACI-7^T^ is considered to represent a novel species of the genus *Methanobacterium* for which the name *Methanobacterium nebraskense* sp. nov. is proposed.

**Table 1.** Differential characteristics of strain ACI-7^T^ and its closest phylogenetic relatives. Strains: 1, ACI-7^T^; 2, *M. oryzae* FPi^T^ (Joulian *et al*., 2000; Schirmack *et al*., 2014); 3, *M. bryantii* M.o.H.^T^ (Bryant *et al*., 1967; Balch *et al*., 1979; Boone *et al*., 2001); 4, *M. veterum* MK4^T^ (Krivushin *et al*., 2010); 5, *M. ivanovii* OCM 140^T^ (Jain *et al*., 1987); 6, *M. movilense* MC-20^T^ (Schirmack *et al*., 2014); 7, *M. uliginosum* P2St^T^ (König 1984; Boone *et al*., 2001); 8, *M. arcticum* M2^T^ (Shcherbakova *et al*., 2011). Cells of all strains are non-motile rods. ND, not determined; +, positive; −, negative.

| Characteristic | 1 | 2 | 3 | 4 | 5 | 6 | 7 | 8 |
| --- | --- | --- | --- | --- | --- | --- | --- | --- |
| Source | Saline wetland soil | Rice field soil | Anaerobic digester | Permafrost sediment | Rock core | Subsurface lake | Marshy soil | Permafrost sediment |
| Cell dimensions (width × length, μm) | 0.2–0.4 × 0.9–4.2 | 0.3–0.4 × 3–10 | 0.5–1.0 × 10–15 | 0.4–0.45 × 2.0–8.0 | 0.5–0.8 × 1.2 | 0.6–0.7 × 3.5–4.0 | 0.2–0.6 × 1.9–3.9 | 0.45–0.5 × 3.0–6.0 |
| Morphology |  |  |  |  |  |  |  |  |
| Chains | + | + | + | + | + | + | + | + |
| Filaments | + | + | + | – | – | – | – | + |
| Aggregates | + | + | + | – | – | – | – | – |
| Temperature range (°C) | 20–45 | 20–42 | ND | 10–46 | ND | 0–44 | 15–45 | 15–45 |
| Optimum temperature (°C) | 40 | 40 | 37–39 | 28 | 45 | 33 | 40 | 37 |
| pH range | 6.5–8.5 | 6.0–8.5 | ND | 5.2–9.4 | 6.5–8.2 | 6.2–9.9 | 6.0–8.5 | 5.5–8.5 |
| Optimum pH | 7.3 | 7.0 | 6.9–7.2 | 7.2–7.4 | 7.0–7.4 | 7.4 | ND | 6.8–7.2 |
| NaCl range (g L<sup>-1</sup>) | 0–25 | 0–25 | ND | 0–17.5 | ND | 0–17.5 | ND | 0–17.5 |
| Substrates |  |  |  |  |  |  |  |  |
| H<sub>2</sub> + CO<sub>2</sub> | + | + | + | + | + | + | + | + |
| Formate | + | + | – | – | – | + | – | + |
| Alcohols | – | – | – | – | – | + | – | – |
| G+C content (mol%)* | 31.8 (G) | 31 (LC) | 33–38 (BD) | 33.8 (T<sub>m</sub>) | 36.6 (T<sub>m</sub>) | 33.0 (LC) | 29.4 (T<sub>m</sub>) | 38.1 (T<sub>m</sub>) |

\*Determined by genome sequencing (G), HPLC (LC), buoyant density (BD), or thermal denaturation (T<sub>m</sub>).

### Description of *Methanobacterium nebraskense* sp. nov

*Methanobacterium nebraskense* (ne.bras.ken′se. N.L. neut. adj. *nebraskense* from Nebraska, where the type strain was isolated).

Cells are non-motile rods, 0.9–4.2 μm in length and 0.2–0.4 µm in diameter, occurring singly, in chains, or as twisted filaments and aggregates. Cells stain Gram-negative and resist lysis by 1% (w/v) SDS and hypotonic solution (distilled water). Colonies on solid medium are small (< 1 mm in diameter) and white with undefined margins, forming within 2 weeks of incubation. Utilizes H_2_ + CO_2_ or formate for growth and methane formation, but not acetate, dimethylsulfide, 2-propanol + CO_2_, methanol, H_2_ + methanol, methylamine, dimethylamine, trimethylamine, or H_2_ + methylamine. Growth factors present in yeast extract are required. Growth occurs from 20–45°C (optimum 40°C), at pH 6.5–8.5 (optimum 7.3), and 0–2.5% NaCl (optimum 0-1%).

The type strain, ACI-7^T^ (=DSM 118696^T^=ATCC TSD-487^T^), was isolated from a saline wetland located in eastern Nebraska, USA. The genomic G+C content of the type strain is 31.79 mol%, as determined by genome sequencing.

## Abbreviations

OGRI: overall genome relatedness index;
ANI: average nucleotide identity;
dDDH: digital DNA–DNA hybridization;
DIC: differential interference contrast;
SDS: sodium dodecyl sulfate;
MES: 2-(*N*-morpholino)ethanesulfonic acid;
PIPES: piperazine-*N*,*N*’-bis(2-ethanesulfonic acid);
HEPES: 4-(2-hydroxyethyl)-1-piperazineethanesulfonic acid;
CHES: *N*-cyclohexyl-2-aminoethanesulfonic acid

## Funding information and Acknowledgements

This work was supported by the Nebraska Center for Energy Sciences Research at the University of Nebraska–Lincoln (to K.A.W.), the National Aeronautics and Space Administration through the NASA Nebraska Space Grant Consortium (Federal Award #80NSSC20M0112 to K.A.W. & N.A.F.), and the National Science Foundation (Graduate Research Fellowship DGE-1610400 to N.A.F. and #2401050 to K.A.W.). Confocal and scanning electron microscopy were carried out at the Microscopy Research Core Facility (RRID, SCR_017798) of the Nebraska Center for Biotechnology at the University of Nebraska–Lincoln, which is partially funded by the Nebraska Center for Integrated Biomolecular Communication (NIH COBRE Award P20-GM113126 and NIGMS) and the Nebraska Research Initiative. Computational analyses were performed using resources provided by the Holland Computing Center of the University of Nebraska, which is supported by the University of Nebraska–Lincoln Office of Research and Innovation and the Nebraska Research Initiative.

