## Supplementary Figures and Tables for "*Methanobacterium nebraskense* sp. nov., a hydrogenotrophic methanogen isolated from saline wetland soil"

**Supplementary Material**

**Fig. S1.** Differential interference contrast and fluorescence microscopy (SYTOX)
**Fig. S2.** Phase contrast and fluorescence microscopy (F420 autofluorescence)
**Fig. S3.** Effect of temperature, pH, and NaCl on methanogenesis

**Table S1.** Accession numbers for taxa in phylogenetic analyses; results for 16S similarity, OGRI
**Table S2.** Genomic evidence for methanogenesis (Wolfe cycle, Wood-Ljungdahl pathway)

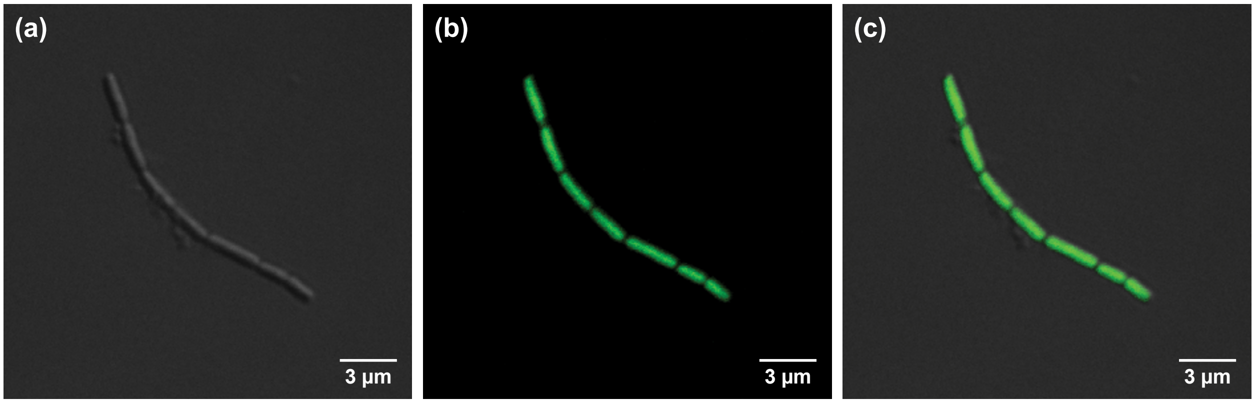


**Fig. S1.** Photomicrographs of strain ACI-7$^T$ obtained with differential interference contrast microscopy (a) and fluorescence microscopy of SYTOX stained cells (b), along with an overlay of both images (c).

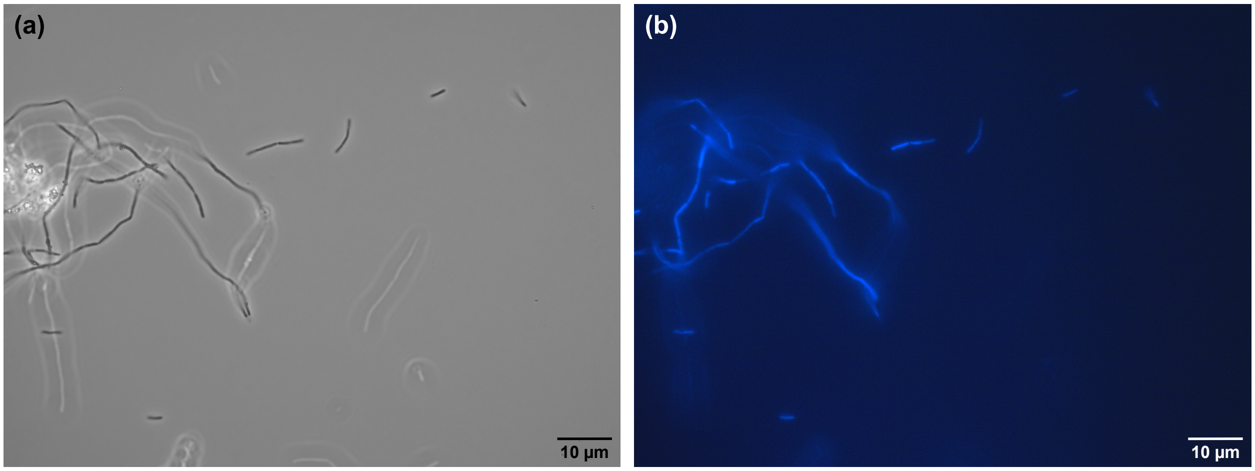


**Fig. S2.** Phase contrast (a) and fluorescence (b) photomicrographs showing $F_{420}$ autofluorescence by cells of strain ACI-7$^T$.

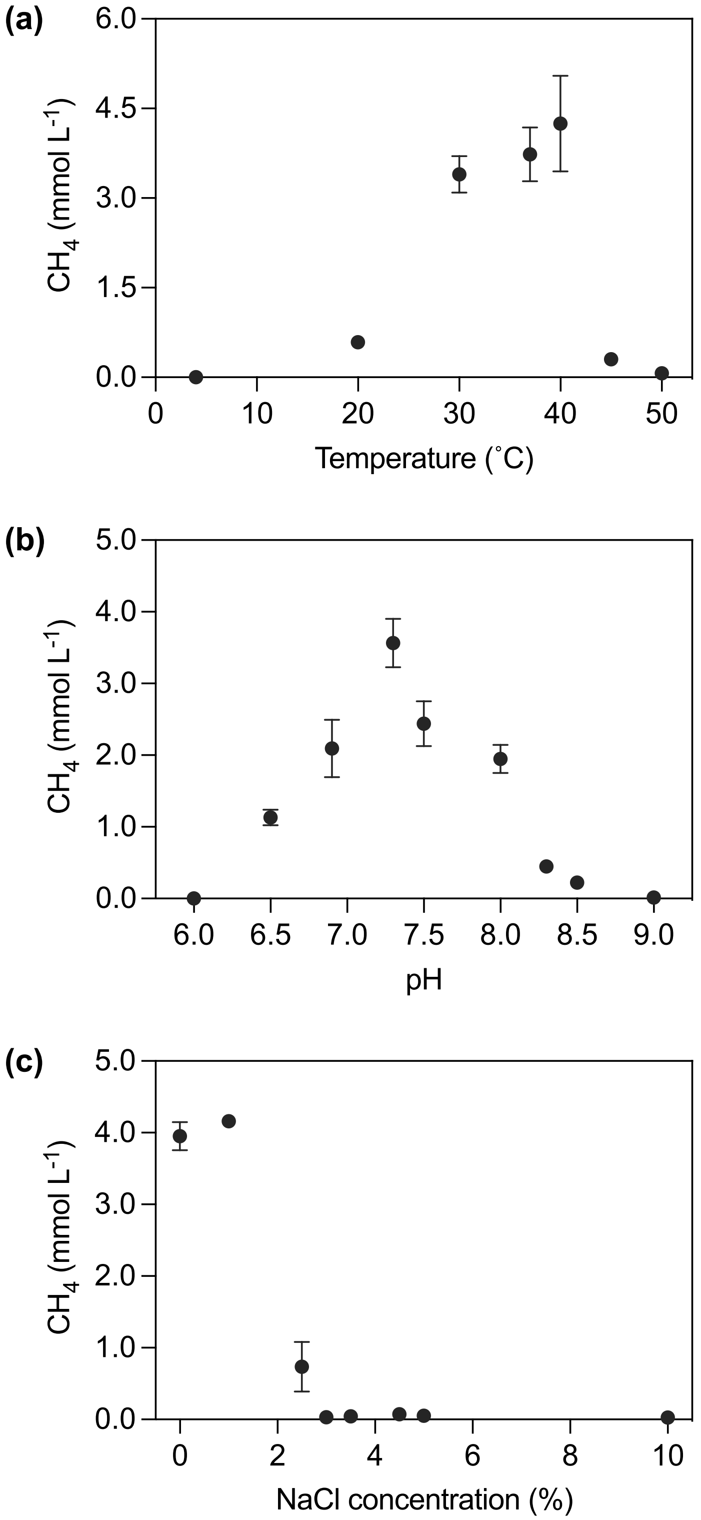


**Fig. S3.** Effect of temperature (a), pH (b), and NaCl concentration (c) on growth of strain ACI-7$^T$. Growth is represented by the total amount of methane accumulated over 5 days (pH) or 7 days (temperature, NaCl) of incubation, in mmol of $CH_4$ per liter of headspace. Error bars represent the standard error of measurement for triplicate cultures; those not visible are smaller than the symbol.

**Table S1.** Strains and accession numbers for *Methanobacterium* species used in phylogenetic and genomic analyses and their respective values for 16S rRNA gene sequence similarity, average nucleotide identity (ANI), and digital DNA–DNA hybridization (dDDH) compared to strain ACI-7$^{T}$. NA, no available genome.

| **Species** | **16S rRNA** | | | **Genome** | | | |
|---|---|---|---|---|---|---|---|
| | **Strain** | **Accession no.** | **16S rRNA similarity (to ACI-7)** | **Strain** | **Accession no.** | **ANI (vs. ACI-7)** | **dDDH (vs. ACI-7)** |
| *Methanobacterium aarhusense* | H2-LR$^{T}$ | NR_042895.1 | 95.27% | – | NA | – | – |
| *Methanobacterium aggregans* | E09F.3$^{T}$ | NR_135896.1 | 95.66% | E09F.3$^{T}$ | GCF_017874455.1 | 70.30% | 20.5% |
| *Methanobacterium alcaliphilum* | WeN4$^{T}$ | NR_028228.1 | 94.17% | WeN3 | GCF_023227715.1 | 70.07% | 19.1% |
| *Methanobacterium alkalithermotolerans* | CAN$^{T}$ | KR349725.1 | 94.78% | CAN$^{T}$ | GCF_018141185.1 | 69.22% | 20.9% |
| *Methanobacterium arcticum* | M2$^{T}$ | NR_115811.1 | 96.47% | M2$^{T}$ | GCF_000746075.1 | 75.56% | 21.1% |
| *Methanobacterium aridiramus* | CWC-01$^{T}$ | MK979366.1 | 94.74% | CWC-01$^{T}$ | GCF_030323845.1 | 68.45% | 20.6% |
| *Methanobacterium beijingense* | 8-2$^{T}$ | NR_028202.1 | 95.05% | – | NA | – | – |
| *Methanobacterium bryantii* | M.o.H.$^{T}$ | NR_042781.1 | 96.68% | M.o.H.$^{T}$ | GCF_002287175.1 | 75.46% | 21.1% |
| *Methanobacterium congolense* | C$^{T}$ | NR_028175.1 | 95.78% | Buetzberg | GCF_900095295.1 | 70.61% | 22.0% |
| *Methanobacterium espanolae* | GP9$^{T}$ | NR_114483.1 | 96.57% | – | NA | – | – |
| *Methanobacterium ferruginis* | Mic6c05$^{T}$ | NR_113045.1 | 94.60% | Mic6c05$^{T}$ | GCF_030296715.1 | 69.66% | 21.4% |
| *Methanobacterium flexile* | GH$^{T}$ | NR_116276.1 | 93.91% | – | NA | – | – |
| *Methanobacterium formicicum* | MF$^{T}$ | NR_115168.1 | 94.64% | Mb9 | GCF_001458655.1 | 69.09% | 20.7% |
| *Methanobacterium ivanovii* | OCM 140$^{T}$ | NR_041716.1 | 96.86% | – | NA | – | – |
| *Methanobacterium kanagiense* | 169$^{T}$ | NR_112749.1 | 94.83% | – | NA | – | – |
| *Methanobacterium lacus* | 17A1$^{T}$ | NR_117917.1 | 95.25% | AL-21 | GCF_000191585.1 | 70.06% | 18.9% |
| *Methanobacterium movens* | TS-2$^{T}$ | NR_116289.1 | 93.92% | – | NA | – | – |
| *Methanobacterium movilense* | MC-20$^{T}$ | NR_133779.1 | 96.73% | – | NA | – | – |
| *Methanobacterium oryzae* | FPi$^{T}$ | NR_028171.1 | 97.09% | FPi$^{T}$ | JGI IMG 2913351602 | 78.17% | 22.0% |
| *Methanobacterium paludis* | SWAN1$^{T}$ | NR_133895.1 | 95.87% | SWAN1$^{T}$ | GCF_000214725.1 | 70.92% | 20.4% |
| *Methanobacterium palustre* | F$^{T}$ | NR_114485.1 | 94.55% | – | NA | – | – |
| *Methanobacterium petrolearium* | Mic5c12$^{T}$ | NR_113044.1 | 95.11% | Mic5c12$^{T}$ | GCA_017873625.1 | 69.21% | 25.7% |
| *Methanobacterium spitsbergense* | VT$^{T}$ | OK037044.1 | 92.78% | VT$^{T}$ | GCF_019931065.1 | 70.76% | 19.6% |
| *Methanobacterium subterraneum* | A8p$^{T}$ | NR_028247.1 | 95.18% | A8p$^{T}$ | GCF_002813695.1 | 69.38% | 21.5% |
| *Methanobacterium thermaggregans* | DSM 3266$^{T}$ | NR_113572.1 | 92.29% | – | NA | – | – |
| *Methanobacterium uliginosum* | P2St$^{T}$ | NR_104694.1 | 96.67% | – | NA | – | – |
| *Methanobacterium veterum* | MK4$^{T}$ | NR_115935.1 | 96.95% | MK4$^{T}$ | GCF_000745485.1 | 75.33% | 21.1% |

**Table S2.** Genes identified in the genome of strain ACI-7$^T$ (accession no. CP166866) associated with the Wolfe cycle and archaeal-type Wood-Ljungdahl pathway and their chromosomal location (in bp, as denoted in the GenBank Record). EC, Enzyme Commission; KO, KEGG Orthology; H4MPT, tetrahydromethanopterin.

| Enzyme | EC # | Gene | KO # | Copies | Location (GenBank Record) |
|---|---|---|---|---|---|
| formylmethanofurandehydrogenase | 1.2.7.12 | fwdA | K00200 | 1 | 382219..383979 |
| | | fwdB | K00201 | 1 | 384810..386108 |
| | | fwdC | K00202 | 1 | 383976..384788 |
| | | fwdD | K00203 | 1 | 381809..382201 |
| | | fwdE | K11261 | 1 | complement(1547157..1547813) |
| | | fwdF | K00205 | 2 | 380451..381563; complement(398349..399359) |
| | | fwdG | K11260 | 1 | 381564..381809 |
| | | fwdH | K00204 | 1 | 379950..380432 |
| formylmethanofuran/H4MPT formyltransferase | 2.3.1.101 | ftr | K00672 | 2 | 1603492..1604367; complement(885025..885918) |
| methenyl-H4MPT cyclohydrolase | 3.5.4.27 | mch | K01499 | 1 | 1348533..1349495 |
| methylene-H4MPT dehydrogenase | 1.5.98.1 | mtd | K00319 | 1 | 639732..640562 |
| methylene-H4MPT reductase | 1.5.98.2 | mer | K00320 | 1 | complement(1146091..1147056) |
| methyl-H4MPT/coenzyme M methyltransferase | 7.2.1.4 | mtrA | K00577 | 1 | 946073..946789 |
| | | mtrB | K00578 | 1 | 945755..946060 |
| | | mtrC | K00579 | 1 | 944921..945742 |
| | | mtrD | K00580 | 1 | 944220..944921 |
| | | mtrE | K00581 | 1 | 943325..944206 |
| | | mtrF | K00582 | 1 | 946802..947008 |
| | | mtrG | K00583 | 1 | 947011..947253 |
| | | mtrH | K00584 | 1 | 947270..948223 |
| methyl-coenzyme M reductase | 2.8.4.1 | mcrA | K00399 | 2 | 941622..943274; 1433188..1434840 |
| | | mcrB | K00401 | 2 | 938459..939787; 1430544..1431875 |
| | | mcrG | K00402 | 2 | 940855..941604; 1432392..1433186 |
| electron-bifurcating hydrogenase-heterodisulfidereductase complex | 1.8.98.5 | mvhA | K14126 | 2 | 50284..51732; 959450..960868 |
| | | mvhD | K14127 | 2 | 810791..811219; 958071..958511 |
| | | mvhG | K14128 | 2 | 49364..50287; 958514..959449 |
| | | hdrA | K03388 | 1 | 518748..520730 |
| | | hdrB | K03389 | 1 | 96387..97322 |
| | | hdrC | K03390 | 1 | 95457..96374 |
| F420-reducing hydrogenase | 1.12.98.1 | frhA | K00440 | 1 | 724520..725743 |
| | | frhB | K00441 | 2 | 137821..138870; 727068..727952 |
| | | frhG | K00443 | 1 | 726229..727056 |
| energy-converting hydrogenase A | | ehaA | K14092 | 1 | 1592657..1592965 |
| | | ehaB | K14093 | 1 | 1592962..1593465 |
| | | ehaC | K14094 | 1 | 1593531..1593779 |
| | | ehaD | K14095 | 1 | 1593798..1594085 |
| | | ehaE | K14096 | 1 | 1594078..1594338 |
| | | ehaF | K14097 | 1 | 1594335..1594856 |
| | | ehaG | K14098 | 1 | 1594811..1595539 |
| | | ehaH | K14099 | 1 | 1595578..1596255 |
| | | ehaI | K14100 | 1 | 1596277..1596492 |
| | | ehaJ | K14101 | 1 | 1596516..1597379 |
| | | ehaK | K14102 | 1 | 1597410..1597667 |
| | | ehaL | K14103 | 1 | 1597678..1598001 |
| | | ehaM | K14104 | 1 | 1597998..1598396 |
| | | ehaN | K14105 | 1 | 1598401..1598850 |
| | | ehaO | K14106 | 1 | 1598847..1599989 |
| | | ehaP | K14107 | 1 | 1600037..1601074 |
| | | ehaQ | K14108 | 1 | 1601071..1602432 |
| | | ehaR | K14109 | 1 | 1602462..1603487 |
| carbon monoxide dehydrogenase/acetyl-CoA synthase | 1.2.7.4; 2.3.1.169; 2.1.1.245 | cdhA | K00192 | 1 | 1093656..1096013 |
| | | cdhB | K00195 | 1 | 1096024..1096533 |
| | | cdhC | K00193 | 1 | 1096579..1097964 |
| | | cdhD | K00194 | 1 | 1098771..1099934 |
| | | cdhE | K00197 | 1 | 1099947..1101320 |
